# The Japan Monkey Centre Primates Brain Imaging Repository of high-resolution postmortem magnetic resonance imaging: the second phase of the archive of digital records

**DOI:** 10.1101/2020.08.23.263517

**Authors:** Tomoko Sakai, Junichi Hata, Yuta Shintaku, Hiroki Ohta, Kazumi Sogabe, Susumu Mori, Hirotaka James Okano, Yuzuru Hamada, Toshiyuki Hirabayashi, Takafumi Minamimoto, Norihiro Sadato, Hideyuki Okano, Kenichi Oishi

## Abstract

A comparison of neuroanatomical features of the brain between humans and our evolutionary relatives, nonhuman primates, is key to understanding the human brain system and the neural basis of mental and neurological disorders. Although most comparative MRI studies of human and nonhuman primate brains have been based on brains of primates that had been used as subjects in experiments, it is essential to investigate various species of nonhuman primates in order to elucidate and interpret the diversity of neuroanatomy features among humans and nonhuman primates. To develop a research platform for this purpose, it is necessary to harmonize the scientific contributions of studies with the standards of animal ethics, animal welfare, and the conservation of brain information for long-term continuation of the field. In previous research, we first developed an open-resource repository of anatomical images obtained using 9.4-T *ex vivo* MRI of postmortem brain samples from 12 nonhuman primate species, and which are stored at the Japan Monkey Centre. In the present study, as a second phase, we released a collection of T2-weighted images and diffusion tensor images obtained in nine species: white-throated capuchin, Bolivian squirrel monkey, stump-tailed macaque, Tibet monkey, Sykes’ monkey, Assamese macaque, pig-tailed macaque, crested macaque, and chimpanzee. Our image repository should facilitate scientific discoveries in the field of comparative neuroscience. This repository can also promote animal ethics and animal welfare in experiments with nonhuman primate models by optimizing methods for *in vivo* and *ex vivo* MRI scanning of brains and supporting veterinary neuroradiological education. In addition, the repository is expected to contribute to conservation, preserving information about the brains of various primates, including endangered species, in a permanent digital form.

## Introduction

The momentum to establish new open data-sharing systems for nonhuman primate neuroimaging with translational possibilities for biomedical research on mental and neurological disorders has recently been increasing (Roelfsema and Treue, 2014) similarly to its human counterpart in the Human Connectome Project (Milham et al., 2020; Milham et al., 2018).

An open-resource repository of nonhuman primate brain images obtained using postmortem (*ex vivo*) MRI, the Japan Monkey Centre (JMC) Primates Brain Imaging Repository, was recently established not only to facilitate scientific discoveries in the field of comparative neuroscience, but also to provide a standard model for animal ethics, animal welfare, and conservation of brain information of various nonhuman primates (http://www.j-monkey.jp/BIR/index_e.html) (Sakai et al., 2018). Following this, we will introduce up-to-date advances in MRI, as well as their limitations, in primate comparative neuroscience, the necessity for consideration of animal ethics, animal welfare, and conservation, and the characteristics of the JMC Primates Brain Sample Collection.

### 1.1. Role of brain MRI in primate comparative neuroscience and current limitations

The comparative study of brain architecture variations among primates, including humans, is absolutely essential for understanding the neurobiological basis of the emergence of the complex brain structure and functions during human evolution. Changes in the brain during evolution from early primates to modern humans involved a marked increase in its overall size, and especially that of the neocortex (Dunbar and Shultz, 2017; Kaas, 2017; Roth and Dicke, 2005), possibly due to genetic or epigenetic changes (Heide et al., 2020; Michael et al.).

Recently, a large-scale MRI study revealed that the traits of the brain system and brain development that are highly phylogenetically conserved show robustness, but the traits that appeared relatively recently during human evolution show large diversity (Reardon et al., 2018). At the same time, some MRI studies of individual differences in human brains suggested that characterizing individual variations in the brain system is critical to understanding the emergence of cognitive and behavioral diversity within populations and may provide insight into the etiology of common neuropsychiatric and neurodevelopmental disorders (Foulkes and Blakemore, 2018; Seghier and Price, 2018).

In addition, interestingly, a recent primate comparative MRI study revealed evidence of evolutionary modifications of human brain connectivity to significantly overlap with the cortical pattern of schizophrenia-related dysconnectivity, suggesting that recent evolutionary modifications in the human brain circuitry may have been one of the factors that played a role in the development of this disorder in humans (van den Heuvel et al., 2019). Therefore, clarifying the cross-species diversity of brain structures among primates is an important issue for primate comparative neuroimaging.

During the past two decades, advances in MRI technology and computational imaging analysis technology have made it possible to conduct direct comparisons of the brain organization of the human with that of nonhuman primates non-destructively. Especially, diffusion MRI is one such noteworthy technology. This imaging technology measures water diffusion as a probe to examine the microscopic organization of anatomical structures within an anatomical region represented by one voxel (Basser et al., 1994a; Basser et al., 1994b). One of the major strengths of diffusion MRI is its capability of delineating well-aligned structures, such as white matter bundles or columnar organization of the cortex. It can also delineate trajectories of major white matter bundles three-dimensionally (Mori and van Zijl, 2002). It has been demonstrated that white matter bundles delineated by diffusion MRI are comparable to those delineated by histology in rhesus macaques (Schmahmann et al., 2007).

In comparative neuroimaging, structural MRI technology has been used to conduct various cross-species comparisons of macrostructures, such as the size of areas and volumes of brain regions (Croxson et al., 2018; Donahue et al., 2018; Eichert et al., 2020; Gomez-Robles et al., 2015; Rilling and Insel, 1999; Rilling and Seligman, 2002; Semendeferi and Damasio, 2000), cortical folding (gyrification) (Amiez et al., 2019; Avants et al., 2006; Glasser et al., 2014; Hopkins et al., 2017; Rilling and Insel, 1999; Van Essen and Dierker, 2007), expansion of cortical areas (Eichert et al., 2020; Hill et al., 2010; Orban et al., 2004; Van Essen and Dierker, 2007), white matter connectivity (Croxson et al., 2018; Reid et al., 2016), and development and aging of the brain (Amlien et al., 2014; Chaplin et al., 2013; Chen et al., 2013; Sakai et al., 2013; Sakai et al., 2011; Sherwood et al., 2011).

Furthermore, *ex vivo* MRI technology has enabled researchers to compare the developmental patterns of gyrification during fetal stages (Hikishima et al., 2013; Sawada et al., 2012; Sawada et al., 2014; Sawada et al., 2009), expansion of the isocortex (Kroenke et al., 2005), and white matter connectivity (Oishi et al., 2011; Rilling et al., 2008) in humans and nonhuman primates.

Recently, many initiatives to provide nonhuman primate MRI databases have been involved in advancing comparative neuroimaging. Details of nonhuman primate neuroimaging resources were reviewed by de Schotten et al. (de Schotten et al., 2019). As a noteworthy database of nonhuman primate MRIs, Milham and colleagues recently launched the Primate Data Exchange (PRIME-DE) initiative, which gathers *in vivo* MRI data from macaques collected by more than 20 different laboratories around the world (The PRIMatE Data Exchange, http://fcon_1000.projects.nitrc.org/indi/indiPRIME.html) (Milham et al., 2020; Milham et al., 2018). In addition, Sherwood, Hopkins, Preuss and colleagues released a chimpanzee brain image database (The National Chimpanzee Brain Resource; NCBR; http://www.chimpanzeebrain.org/). Also, the Toro and Heuer group has collected structural data from postmortem samples of nonhuman primates and other animals (Brain Catalogue; https://braincatalogue.org) (Heuer et al., 2019).

The new open science approach for providing nonhuman primate MRI resources enables researchers to collect detailed information about brain anatomy to facilitate their comparative neuroimaging studies, but they nevertheless face some inevitable limitations.

First, to date, most studies using the nonhuman primate MRI data resource described above have focused solely on experimental primate models such as macaque monkeys (de Schotten et al., 2019). To better understand the brain’s anatomical features and connectivity across primate lineages, more species will need to be comprehensively studied.

In this regard, the Toro and Heuer group published *ex vivo* structural MRI data from 31 nonhuman primate species (Heuer et al., 2019). However, their scanning methods lacked uniformity in terms of imaging techniques (different devices and imaging sequences were used) and postmortem samples (a mixture of extracted brain samples, brain samples including skulls, and whole-body samples was used). Although they quantified the quality of those data, those limitations may handicap any meaningful comparative analyses of nonhuman primate brains. In order to perform a more robust comparative analysis, it may be necessary to image extracted brains that have undergone the same pretreatment procedures using the identical high-field MRI equipment.

Finally, most of the previous image databases of nonhuman primate brains show few MRIs obtained by the use of 7-T or higher-field (ultra-high field) MRI scanners. In fact, it has been difficult to collect high-resolution diffusion tensor image (DTI) data for nonhuman primate brain samples larger than marmoset brain specimens.

### 1.2. Necessary consideration of animal ethics, welfare, and conservation

It is important for researchers who study nonhuman primates to contribute not only to their own study but also to animal welfare and ethics (Amadio et al., 2018; Buller, 2018; Milham et al., 2020; Milham et al., 2018; Sadato et al., 2019). The challenge of understanding the human brain and its disorders means that nonhuman primate research is increasingly needed, although it is essential that this research is complemented by studies using other approaches, and other alternatives to (*in vivo*) nonhuman primate use (Lemon, 2018). Application of the 3Rs (Replacement, Reduction, Refinement) (Russell and Burch, 2009; Russell and Burch, 1959) has been led by the UK National Centre for the 3Rs (NC3Rs), with active participation of nonhuman primate researchers, who are constantly refining the research methodology used around the world.

Sharing large-scale digital information and data obtained by the postmortem brain MRI approach with large numbers of researchers contributes to the elimination of duplicate nonhuman primate experiments and minimizes the number of animals used for scientific purposes. Moreover, elucidating spontaneously occurring brain disorders of nonhuman primates using an *ex vivo* MRI approach can be of great significance, as a kind of “autopsy imaging of nonhuman primates”.

The digital brain records obtained by the *ex vivo* MRI approach are advantageous for permanently archiving information about endangered species. It has been reported that about 60% of nonhuman primate species are now threatened with extinction. Not only members of the hominoid family, such as gibbons, orangutans (*Pongo*), gorillas, and chimpanzees, but also all 16 extant primate families are facing extinction (Estrada et al., 2017). Further, the populations of 75% of primate species are decreasing globally (Estrada et al., 2017). This highlights the significance of recording and storing information about these species permanently in electronic form.

### 1.3. The Japan Monkey Centre primate postmortem brain collection

The JMC, one of the largest nonhuman primate museums and zoos in the world, was founded at Inuyama, Japan, in 1956, with the support of Nagoya Railroad Co., Ltd. In 2014, the JMC was transformed into a public interest foundation. The JMC aims to contribute to the development of science, education, culture, and the harmony of local communities and the global environment on the basis of research, conservation, social education, publication, management of its zoo, and the collection of specimens and other materials regarding nonhuman primates.

At present, 850 nonhuman primates of approximately 60 species live at the JMC in an area of 43,600 m^2^. In terms of animal ethics and welfare, consideration must be given to not only the physical but also the psychological well-being of captive nonhuman primates in research settings and zoos (Lutz and Novak, 2005). In recent years, similarly to modern zoos, JMC has been paying heed to animal welfare; for example, (1) captive nonhuman primates are housed in groups so that they can maintain the societies of their respective species, (2) three-dimensional structures and playground equipment have been introduced as part of the expansion of available space for captive nonhuman primates, (3) varieties of food items and feeders are provided to enhance the feeding experience (Fig.1). In particular, it must be noted that the JMC postmortem brain samples were obtained from individuals that had extensive species-specific social experiences in a socially enriched environment. This advantage would be difficult to fully realize in other typical animal experimental facilities. The JMC’s social enrichment environment allows us to make a direct cross-species comparison of the relationship between brain structures and social features (such as allomothering and social structure) among primates.

**Fig.1.**
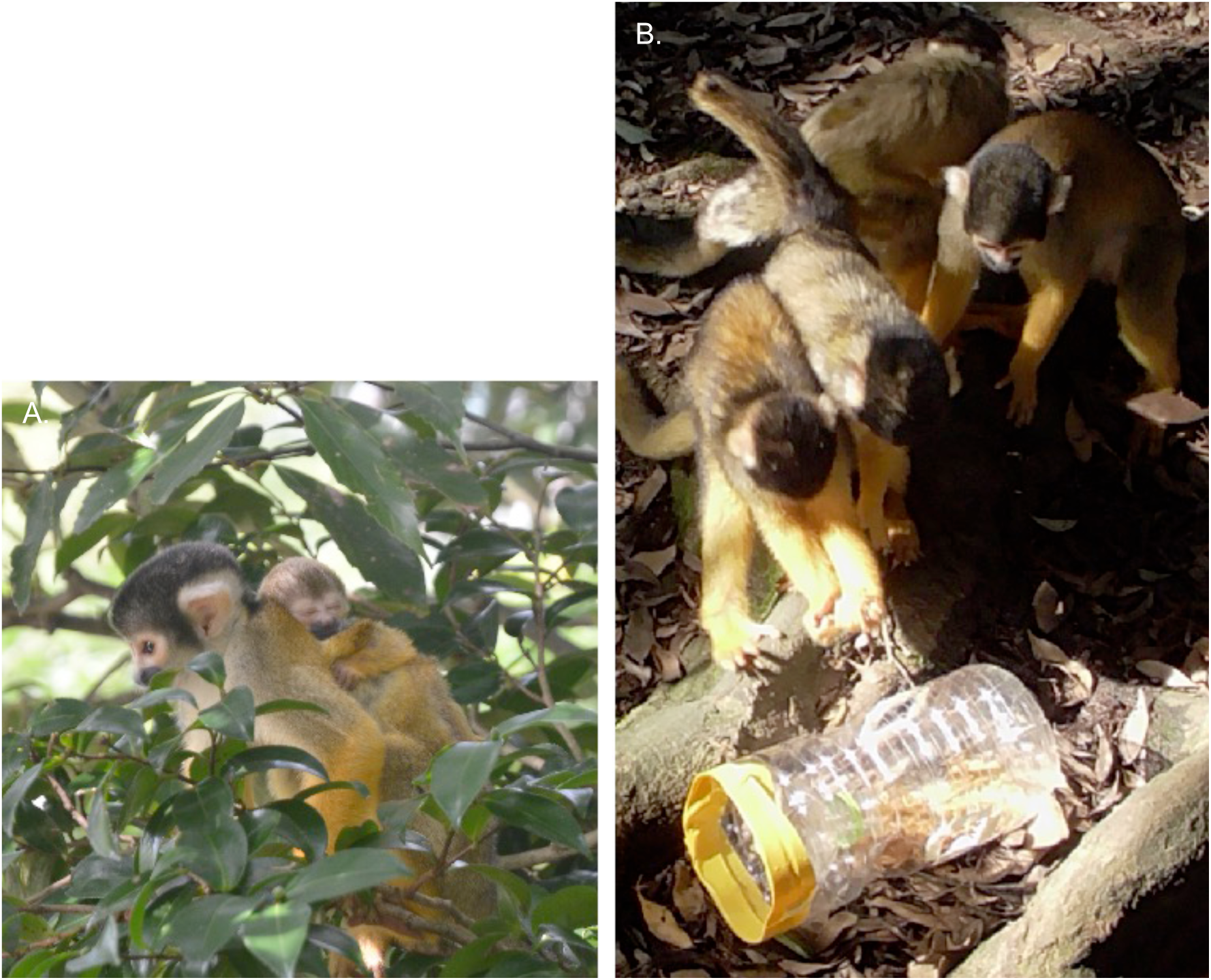
Animal welfare considerations for nonhuman primates at the JMC. Various considerations are made for captive nonhuman primates from the viewpoint of animal welfare at the JMC. (A) Mother and child Bolivian squirrel monkeys living in an enriched social and physical environment. (B) Bolivian squirrel monkeys looking for foods in a food box made from plastic bottles.

Since the foundation of the JMC in 1956, more than 2400 postmortem brain samples have been collected from the individuals from more than 100 nonhuman primate species from 11 families, including endangered species such as gibbons (*Hylobatidae*), gorillas (*Gorilla*), chimpanzees (*Pan troglodytes)*, and vulnerable species such as owl monkeys (*Aotus trivirgatus*). Details of the JMC Primate Brain Sample Collection are listed in Table 1. These brain samples were prepared by skilled veterinarians, who extracted brains from the bodies of nonhuman primates that had died spontaneously. In summary, the rearing of nonhuman primates and collecting their postmortem brain samples at the JMC can be said to be fully considered according to animal ethics and welfare.

**Table 1.** The JMC Primate Brain Sample Collection

| Families | Number of species | Number of brain samples |
| --- | --- | --- |
| Cheirogaleidae | 1 | 7 |
| Lemuridae | 4 | 55 |
| Galagidae | 2 | 25 |
| Lorisidae | 5 | 56 |
| Tarsiidae | 1 | 5 |
| Cebidae | 26 | 693 |
| Atelidae | 8 | 61 |
| Pitheciidae | 6 | 32 |
| Cercopithecidae | 55 | 1470 |
| Hylobatidae | 6 | 39 |
| Hominidae | 4 | 19 |

(Total 11 Families, 118 species, 2462 brain samples as of April 2020)

### 1.4. First phase of Japan Monkey Centre Primates Brain Imaging Repository and its limitations

To accelerate research in the field of comparative neuroscience, and to contribute to animal ethics, welfare, and conservation, we launched a project to create a brain MRI repository consisting of MRI data of the brains of various nonhuman primates. As the initial phase of creating an open resource, we developed a collection of structural MRIs and diffusion tensor images (DTIs) of brains obtained from 12 species, stored in the JMC, that were scanned using a 9.4-T MRI scanner: pygmy marmoset (*Cebuella pygmaea*), owl monkey, white-fronted capuchin (*Cebus albifrons*), crab-eating macaque (*Macaca fascicularis*), Japanese macaque (*Macaca fuscata*), bonnet macaque (*Macaca radiata*), toque macaque (*Macaca sinica*), Sykes’ monkey (*Cercopithecus albogularis*), red-tailed monkey (*Cercopithecus ascanius*), Schmidt’s guenon (*Cercopithecus ascanius schmidti*), de Brazza’s guenon (*Cercopithecus neglectus*), and lar gibbon (*Hylobates lar*) (The JMC Brain Imaging Repository, http://www.j-monkey.jp/BIR/index_e.html) (Sakai et al., 2018).

However, this first phase of the development had several technical limitations regarding the postmortem MRI. A more detailed explanation of these technical limitations is presented in “4.1. Technical improvements from first phase in high-resolution postmortem MRI” in the Discussion section.

### 1.5. Goals of second phase of Japan Monkey Centre Primates Brain Imaging Repository

As the second phase, by solving the above described technical limitations, we acquired structural MRI and DTI data from nine nonhuman primate species using the same high-resolution MRI: white-throated capuchin (*Cebus capucinus*), Bolivian squirrel monkey (*Saimiri boliviensis*), stump-tailed macaque (*Macaca arctoides*), Tibet monkey (*Macaca thibetana*), Sykes’ monkey, Assamese macaque (*Macaca assamensis*), pig-tailed macaque (*Macaca nemestrina*), crested macaque (*Macaca nigra*), and chimpanzee.

In other words, the data were acquired from two species from the family Cebidae: white-throated capuchin and Bolivian squirrel monkey, six species from the family Cercopithecidae: stump-tailed macaque, Tibet monkey, Sykes’ monkey, Assamese macaque, pig-tailed macaque, crested macaque, and one species from the Hominidae: chimpanzee.

Brain samples from eight (white-throated capuchin, Bolivian squirrel monkey, stump-tailed macaque, Tibet monkey, Assamese macaque, pig-tailed macaque, crested macaque, chimpanzee) of the nine nonhuman primate species were the first to be scanned in the second phase of the JMC Primates Brain Imaging Repository. The brain samples from six (Bolivian squirrel monkey, stump-tailed macaque, Tibet monkey, Assamese macaque, pig-tailed macaque, crested macaque) of the nine species were the first brain samples of these species in the world to be successfully scanned by MRI. Then, we added these data to update the JMC Primates Brain Imaging Repository in order to make the electronical image data available to the science community.

## 2. Methods

### 2.1. JMC primate postmortem brain collection

Twelve postmortem brain samples from nine species were selected from the JMC primate postmortem collection in this study (Table 2). Brain samples were immersed in 10% neutral buffered formalin and fixed for more than 1 month. All procedures described herein were performed in accordance with a protocol approved by the JMC’s review board for primate research (permits #2014013, #2015019, and #2016017).

**Table 2.**
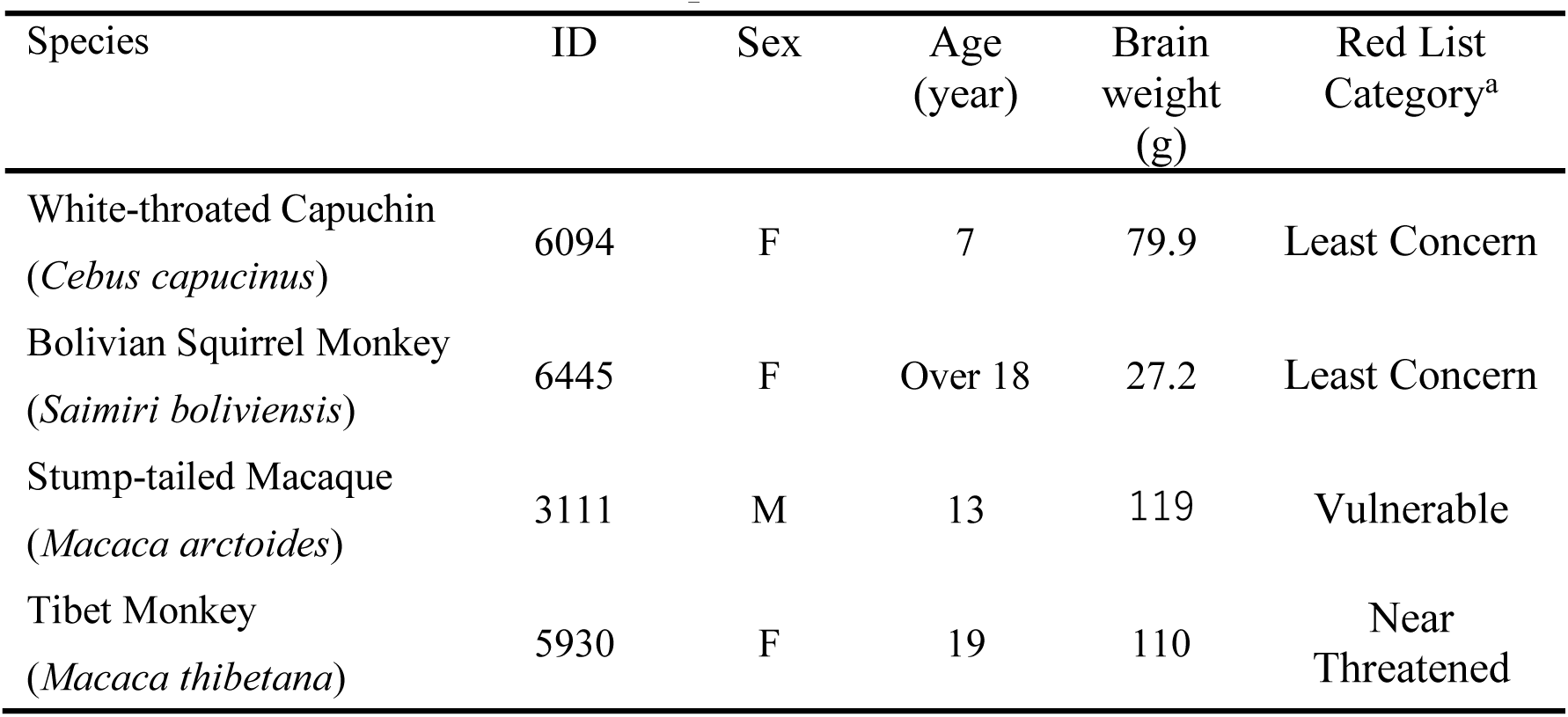
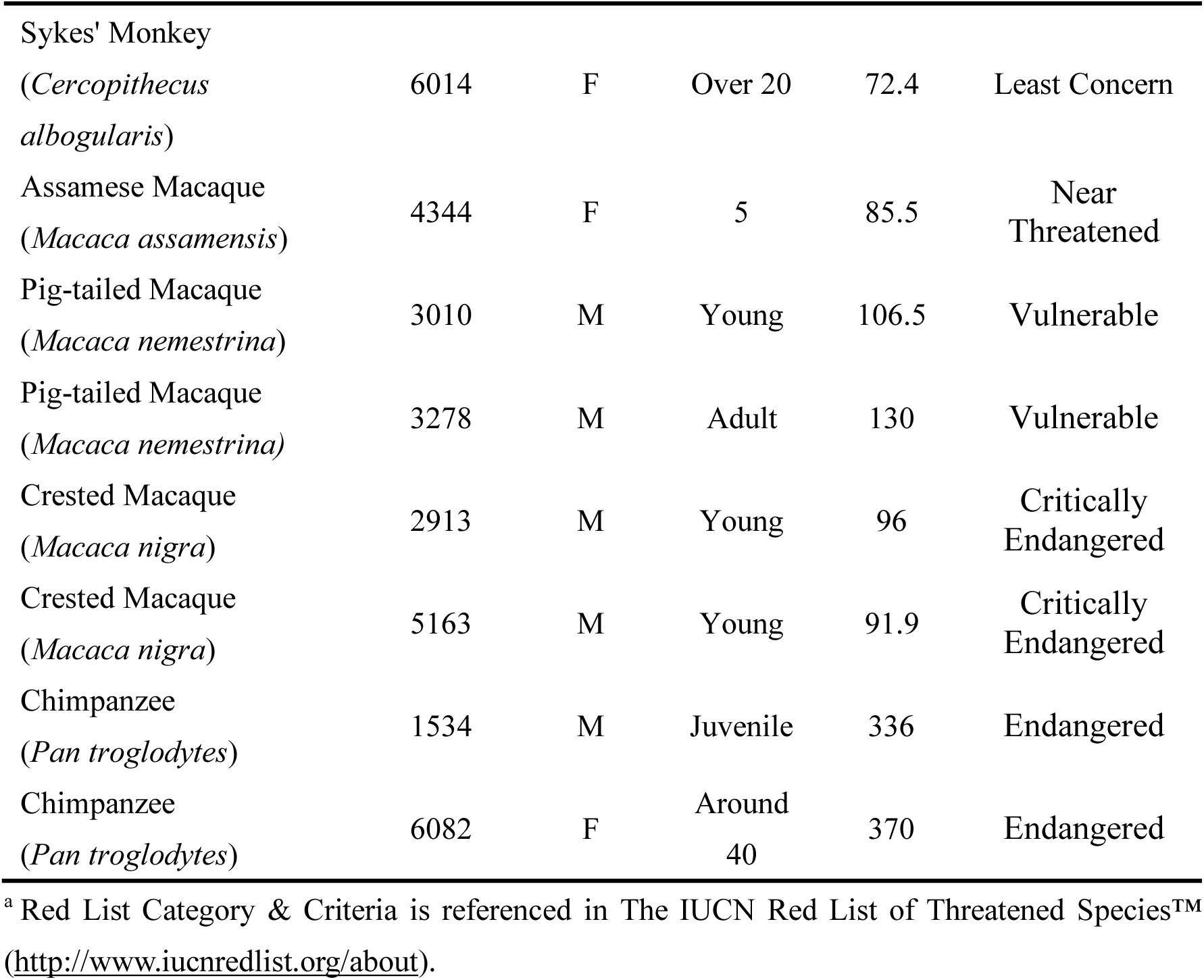
Characteristics of brain samples

| Species | ID | Sex | Age<br>(year) | Brain<br>weight<br>(g) | Red List<br>Category<sup>a</sup> |
| --- | --- | --- | --- | --- | --- |
| White-throated Capuchin<br>(*Cebus capucinus*) | 6094 | F | 7 | 79.9 | Least Concern |
| Bolivian Squirrel Monkey<br>(*Saimiri boliviensis*) | 6445 | F | Over 18 | 27.2 | Least Concern |
| Stump-tailed Macaque<br>(*Macaca arctoides*) | 3111 | M | 13 | 119 | Vulnerable |
| Tibet Monkey<br>(*Macaca thibetana*) | 5930 | F | 19 | 110 | Near<br>Threatened |
| Sykes' Monkey<br>(*Cercopithecus albogularis*) | 6014 | F | Over 20 | 72.4 | Least Concern |
| Assamese Macaque<br>(*Macaca assamensis*) | 4344 | F | 5 | 85.5 | Near Threatened |
| Pig-tailed Macaque<br>(*Macaca nemestrina*) | 3010 | M | Young | 106.5 | Vulnerable |
| Pig-tailed Macaque<br>(*Macaca nemestrina*) | 3278 | M | Adult | 130 | Vulnerable |
| Crested Macaque<br>(*Macaca nigra*) | 2913 | M | Young | 96 | Critically Endangered |
| Crested Macaque<br>(*Macaca nigra*) | 5163 | M | Young | 91.9 | Critically Endangered |
| Chimpanzee<br>(*Pan troglodytes*) | 1534 | M | Juvenile | 336 | Endangered |
| Chimpanzee<br>(*Pan troglodytes*) | 6082 | F | Around 40 | 370 | Endangered |

<sup>a</sup> Red List Category & Criteria is referenced in The IUCN Red List of Threatened Species™ (http://www.iucnredlist.org/about).

### 2.2. Preparation for postmortem MRI

We prepared a workstation with a brain sample, small sponges, a polyethylene container customized for the brain samples, Fluorinert™ (3M Japan Limited, Tokyo, Japan), and a vacuum pump. The preparation procedure is as follows: (1) completely remove water from containers and sponges; (2) fill the bottom of the container with the sponges; (3) gently dry formalin from the surface of the brain sample with a paper towel; (4) insert the brain sample with the frontal pole toward the bottom of the container; (5) fill the container with Fluorinert™ to approximately 80% of the container volume to prevent dehydration and reduce susceptibility to artifacts at tissue margins; (6) carefully secure the brain sample in the container using more sponges around the sides to fix its position; (7) fill the rest of the container with sponges and Fluorinert™; (8) place the container in a chamber connected to a vacuum pump and deaerate the contents of the container until no air bubbles are trapped in interstices of the sulci; although it depends on the size of the brain sample, the deaeration takes approximately 60 minutes on average; if the brain sample itself is sufficiently deaerated, some air remaining in the sponges is inconsequential; (9) secure the cap and seal the container with a plastic tape; (10) insert the container into the MRI coil; maintain the temperature of the brain samples at 24°C during the scans in accordance with the International Industrial Standard (IEC 60601-2-33) (Bottomley et al., 1984; Fix et al., 2000) (Fig. 2).

**Fig.2.**
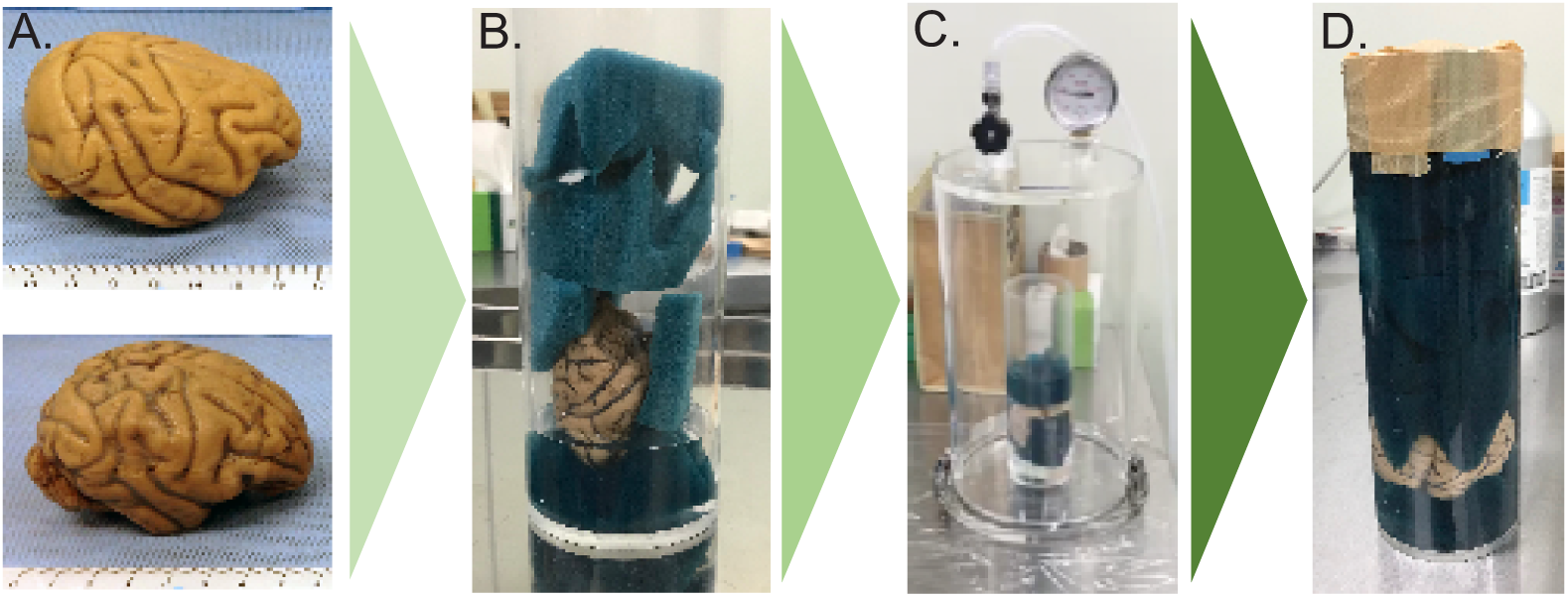
Preparation for postmortem MRI. Outline of the preparation procedure for postmortem. (A) Gently dry/soak up the formalin from the brain sample surface with a paper towel. (B) Samples are placed in a customized polyethylene container filled with Fluorinert™ to prevent dehydration and reduce susceptibility to artifacts at tissue margins. (C) The container is placed in a vacuum pump and the contents of the container are deaerated until no air bubbles are trapped in interstices of sulci. (D) Cap is secured and container is sealed with plastic tape.

### 2.3. Image acquisition

In the second phase, by using 9.4-T MRI equipment with a larger bore, we attempted to scan whole brain samples of great apes that could not be scanned in the first phase. The detailed scanning protocols were as follows. The whole-brain scans for chimpanzee brain samples were performed using a 9.4-T Biospec 90/30 MRI scanner (Bruker Biospin GmbH; Ettlingen, Germany) with a 154-mm inner diameter. The whole-brain scans for the other nonhuman primate brain samples were performed using a 9.4-T Biospec 90/20 MRI scanner (Bruker Biospin GmbH; Ettlingen, Germany) with a triple-axis gradient system (maximum gradient strength of 300 mT/m), using a transmitting and receiving two-channel quadrature coil with an 86-mm inner diameter. T2-weighted images were acquired using a three-dimensional rapid acquisition with refocused echoes (RARE) sequence. DTIs were acquired using a conventional pulsed gradient spin echo (PGSE) sequence. Acquisition parameters and lengths of scan time for T2-weighted images and DTIs are shown in Tables 3 and 4, respectively.

**Table 3.** Brain samples evaluated by T2-weighted image

| Species | ID | TE, TR,<br>NEX<sup>a</sup> | FOV<sup>b</sup><br>(mm<sup>3</sup>) | Matrix size | Isotropic<br>spatial<br>resolution<br>(mm) | Scan<br>time<sup>c</sup> |
| --- | --- | --- | --- | --- | --- | --- |
| White-throated Capuchin | 6094 | 7.5, 600, 3 | 64, 51.2, 43.2 | 320, 256, 216 | 0.2 | 7h29m |
| Bolivian Squirrel Monkey | 6445 | 7.5, 600, 3 | 52.2, 40.8, 33 | 348, 272, 220 | 0.15 | 5h12m |
| Stump-tailed Macaque | 3111 | 7.5, 600, 2 | 80, 64, 48 | 320, 256, 192 | 0.25 | 4h28m |
| Tibet Monkey | 5930 | 7.5, 600, 3 | 84, 67.2, 50.4 | 320, 256, 192 | 0.21 | 6h43m |
| Sykes' Monkey | 6014 | 7.5, 600, 4 | 73.6, 58.9, 44.2 | 320, 256, 192 | 0.23 | 8h57m |
| Assamese Macaque | 4344 | 7.5, 600, 4 | 72, 57.6, 43.2 | 320, 256, 192 | 0.23 | 8h57m |
| Pig-tailed Macaque | 3010 | 7.5, 600, 3 | 72, 57.6, 48.6 | 320, 256, 216 | 0.23 | 7h33m |
| Pig-tailed Macaque | 3278 | 7.5, 600, 3 | 80, 64, 54 | 320, 256, 216 | 0.25 | 7h33m |
| Crested Macaque | 2913 | 7.5, 600, 4 | 72, 57.6, 43.2 | 320, 256, 192 | 0.23 | 8h57m |
| Crested Macaque | 5163 | 7.5, 600, 4 | 76.8, 61.4, 46 | 320, 256, 192 | 0.24 | 8h57m |
| Chimpanzee | 1534 | 15, 400, 2 | 125, 100, 80 | 500, 400, 320 | 0.25 | 7h6m |
| Chimpanzee | 6082 | 15, 400, 2 | 125, 100, 80 | 500, 400, 320 | 0.25 | 7h6m |

<sup>a</sup> Echo time (TE) and repetition time (TR) are in units of seconds. NEX is the number of excitations used for MR signal averaging.
<sup>b</sup> Field of view (FOV)
<sup>c</sup> Scan times are in units of hours (h) and minutes (m)

**Table 4.** Brain samples evaluated by DTI

| Species | ID | TE, TR,<br>NEX<sup>a</sup> | FOV<sup>b</sup><br>(mm<sup>3</sup>) | Matrix<br>size | Isotropic<br>spatial<br>resolution<br>(mm) | b0<br>im<br>age | Scan<br>time<sup>c</sup> |
| --- | --- | --- | --- | --- | --- | --- | --- |
| White-throated Capuchin | 6094 | 20, 650, 1 | 64, 51.2, 43.2 | 160, 128, 108 | 0.4 | 2 | 59h54m |
| Bolivian Squirrel Monkey | 6445 | 20.1, 630, 1 | 52.2, 40.8, 33 | 174, 136, 110 | 0.3 | 2 | 62h49m |
| Stump-tailed Macaque | 3111 | 20, 650, 1 | 80, 64, 48 | 160, 128, 96 | 0.5 | 2 | 53h14m |
| Tibet Monkey | 5930 | 20, 650, 1 | 84, 67.2, 50.4 | 160, 128, 96 | 0.53 | 2 | 53h14m |
| Sykes' Monkey | 6014 | 20, 700, 1 | 73.6, 58.9, 44.2 | 160, 128, 96 | 0.46 | 3 | 59h8m |
| Assamese Macaque | 4344 | 20,700, 1 | 72, 57.6, 43.2 | 160, 128, 96 | 0.45 | 2 | 57h20m |
| Pig-tailed Macaque | 3010 | 20, 650, 1 | 72, 57.6, 48.6 | 160, 128, 108 | 0.45 | 2 | 59h54m |
| Pig-tailed Macaque | 3278 | 20, 600, 1 | 80, 64, 54 | 160, 128, 108 | 0.5 | 2 | 55h17m |
| Crested Macaque | 2913 | 20, 700, 1 | 72, 57.6, 43.2 | 160, 128, 96 | 0.45 | 2 | 57h20m |
| Crested Macaque | 5163 | 20, 650, 1 | 76.8, 61.4, 46 | 160, 128, 96 | 0.48 | 2 | 53h14m |
| Chimpanzee | 1534 | 28.9, 300, 1 | 125, 100, 80 | 250, 200, 160 | 0.5 | 2 | 76h22m |
| Chimpanzee | 6082 | 28.9, 300, 1 | 125, 100, 80 | 250, 200, 160 | 0.5 | 2 | 77h39m |

<sup>a</sup> Echo time (TE) and repetition time (TR) are in units of seconds. NEX is the number of excitations used for MR signal averaging.
<sup>b</sup> Field of view (FOV)
<sup>c</sup> Scan times are in units of hours (h) and minutes (m)

### 2.4. Image processing

T2-weighted images were processed following a series of procedures using ANALYZE v. 9.0 software (Mayo Clinic, Mayo Foundation, Rochester, MN, USA) and ROIEditor (https://www.mristudio.org) (Jiang et al., 2006), and DiffeoMap (https://www.mristudio.org) (Fig. 3). (1) T2-weighted images were resampled to isotropic voxels using ANALYZE v. 9.0. Image resolution varied depending on the species, ranging from 0.15 to 0.25 mm. (2) The resampled T2-weighted image was in a standard anatomical orientation, with the transaxial plane parallel to the anterior commissure-posterior commissure (AC-PC) line and perpendicular to the interhemispheric fissure on ROIEditor. (3) A radio-frequency bias field correction was applied using the Bias Correction function implemented in DiffeoMap.

**Fig.3.**
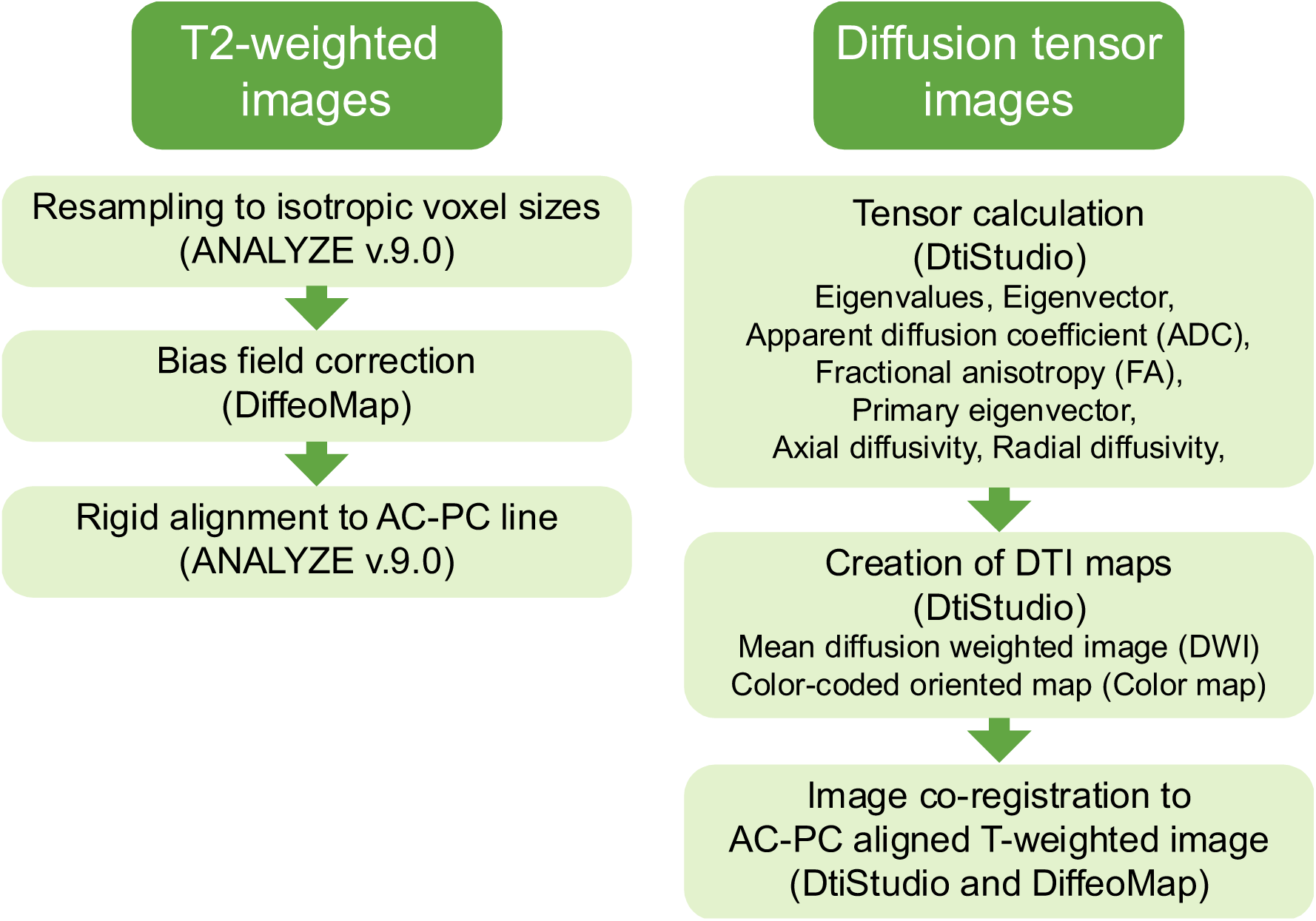
Schema of image processing. T2-weighted images were first processed by resampling to isotropic voxel size, followed by bias field correction and rigid alignment to AC-PC line. Diffusion tensor images were then processed by tensor calculation, followed by map creation. Finally, the diffusion tensor images were co-registered to the AC-PC-aligned T2-weighted images.

DTIs of each brain were analyzed using the following series of procedures using DtiStudio (https://www.mristudio.org) (Jiang et al., 2006; Mori and Zhang, 2006) and DiffeoMap (https://www.mristudio.org). (1) The eigenvalues and associated eigenvectors (Basser et al., 1994b) of the voxel-wise diffusion tensors were computed at each pixel along with the apparent diffusion coefficient (ADC), fractional anisotropy (FA), primary eigenvector, axial diffusivity (λ|, the primary eigenvalue), and radial diffusivity (λ⊥, average of the secondary and tertiary eigenvalues). (2) The mean diffusion-weighted image (DWI) was generated by averaging all diffusion-weighted images. (3) The color-coded orientation map (color map) was created by assigning red, green, and blue components equal to the ratio of the absolute values of x (medial-lateral), y (anterior-posterior) and z (superior-inferior) components of the primary eigenvector, where intensity was proportional to FA (Makris et al., 1997; Pajevic and Pierpaoli, 1999). (4) DWI, FA, and color maps for each sample were resliced to the isotropic voxel identical to that of the corresponding T2-weighted image on DiffeoMap. (5) The least diffusion-weighted image (b0) of the DTI was transformed to the T2-weighted image using six-parameter rigid transformation of the Automatized Image Registration (AIR) (Woods et al., 1998) implemented on DiffeoMap, and the transformation was applied to the corresponding DWI and tensor field, from which FA and color maps were recalculated. (6) All images were saved in the ANALYZE format.

The T2-weighted images and DTIs were quality-controlled by the authors according to the criteria of previous studies (Sakai et al., 2018), and those with poor quality were excluded from the repository. Diffusion toolkit and Track Viz (http://trackvis.org) (Wang et al., 2007) were used to reconstruct and visualize tractography (Mori et al., 1999).

## 3. Results

We acquired high-spatial-resolution T2-weighted images and DTIs (ex, DWIs, b0, FA map, color map and tractography) from various nonhuman primate species (Figs. 4–11). We excluded a set of DTIs from the Stump-tailed macaque (Pr5930) from the original datasets collected, due to extensive image error. Thus, we showed T2-weighted images from 12 individuals belonging to nine species (Fig.4) and DTIs from 11 individuals belonging to nine species (Fig.5). In this study, we identified the common sulci, areas, and white matter bundles of the brain among primate species to present the quality of the acquired brain images. We used a heterological atlas of the rhesus macaque brain (Paxinos et al., 2000) and a DTI atlas of the human brain (Mori et al., 2005; Oishi et al., 2010) to identify the anatomical features and white matter bundles. Detailed results were as below.

**Fig.4.**
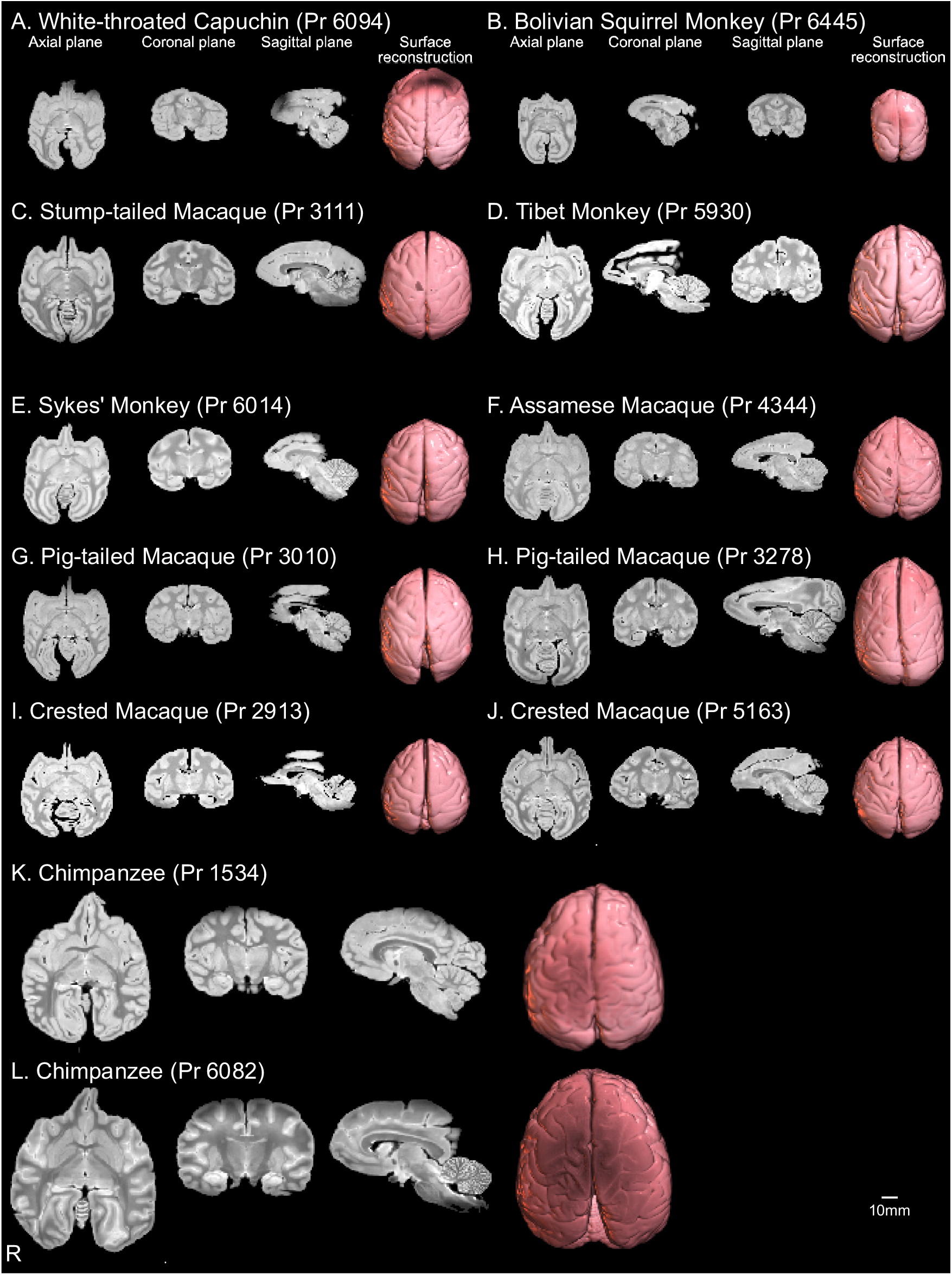
Representative samples of T2-weighted images and brain surfaces of 9 non-human primate brain samples in the database. T2-weighted images and surface reconstructions of the brain for each brain sample in these 9 nonhuman primate species: (A) White-throated Capuchin (Pr 6094), (B) Bolivian Squirrel Monkey (Pr 6445), (C) Stump-tailed Macaque (Pr 3111), (D) Tibet Monkey (Pr 5930), (E) Sykes’ Monkey (Pr 6014), (F) Assamese Macaque (Pr 4344), (G) Pig-tailed Macaque (Pr 3010), (H) Pig-tailed Macaque (Pr 3278), (I) Crested Macaque (Pr 2913), Crested Macaque (Pr 5163), (I) Chimpanzee (Pr 1534), Chimpanzee (Pr 6082).

**Fig.5.**
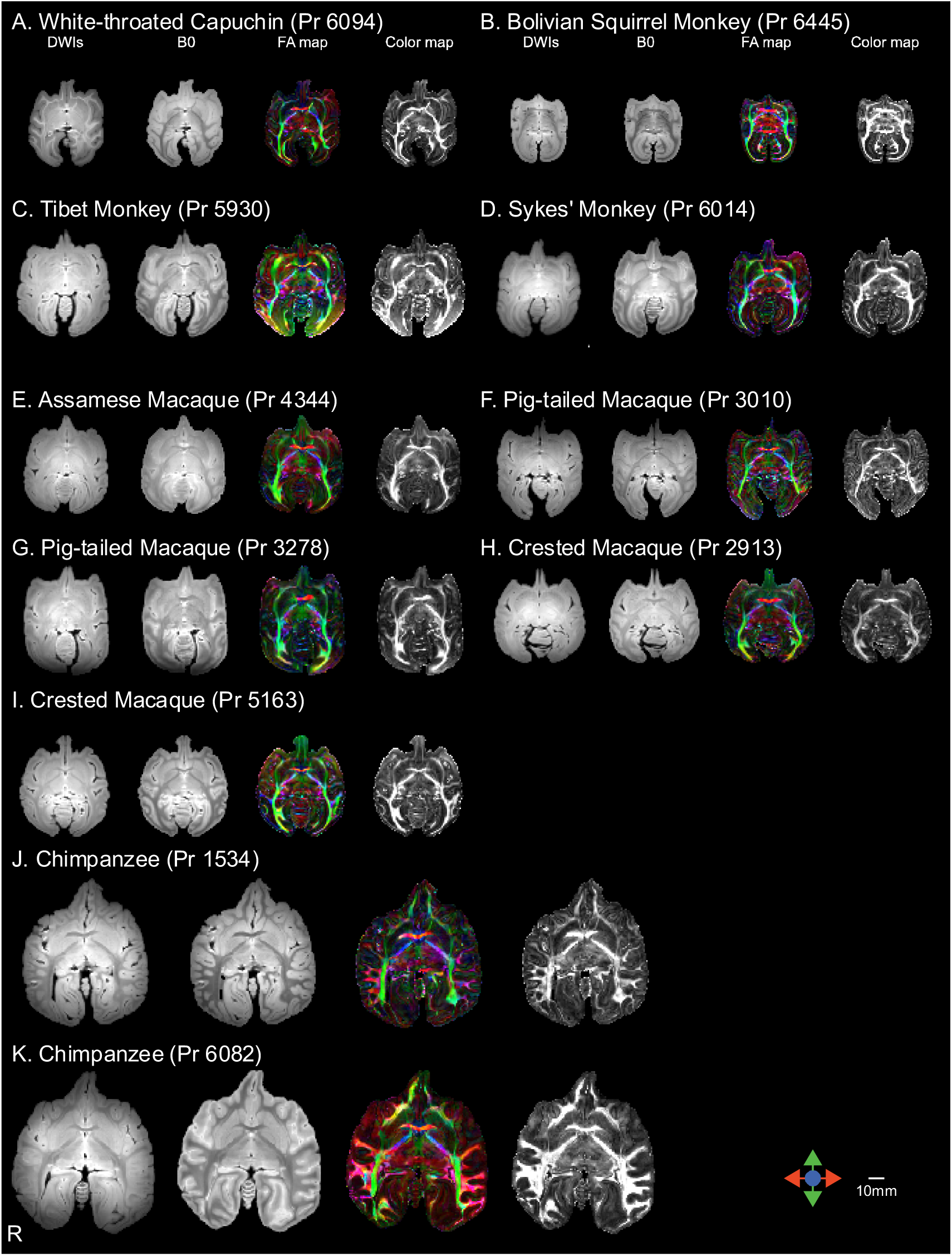
Representative samples of DTI images of 11 non-human primate brain samples in the database. The images are DWIs, b0, FA and Color maps from the mid-axial plane for each brain sample in these 9 nonhuman primate species: (A) White-throated Capuchin (Pr 6094), (B) Bolivian Squirrel Monkey (Pr 6445), (C) Tibet Monkey (Pr 5930), (D) Sykes’ Monkey (Pr 6014), (E) Assamese Macaque (Pr 4344), (F) Pig-tailed Macaque (Pr 3010), (G) Pig-tailed Macaque (Pr 3278), (H) Crested Macaque (Pr 2913), (I) Crested Macaque (Pr 5163), (J) Chimpanzee (Pr 1534), (K) Chimpanzee (Pr 6082).

### 3.1. Anatomical features from T2-weighted images

Representative results of the T2-weighted images from nine species are shown in Fig. 4. The sulci and gyri were identified in the brain surface of T2-weighted images (Fig. 6). For example, the lateral view of the chimpanzee (Pr1534) brain displayed the infraprincipal dimple, inferior arcuate sulcus, central sulcus, intraparietal sulcus, lateral fissure, superior temporal sulcus, lunate sulcus, inferior occipital sulcus, and external calcarine sulcus (Fig. 6). The middle temporal gyrus and temporal polar gyrus were also delineated. In the dorsal view of the brain showed the anterior supraprincipal dimple, superior arcuate sulcus, superior precentral dimple, central sulcus, intraparietal sulcus, superior postcentral dimple, superior temporal sulcus, and parieto-occipital sulcus (Fig. 6). The superior frontal gyrus, middle frontal gyrus, inferior frontal gyrus, precentral gyrus, anterior superior parietal gyrus, cingulate gyrus, posterior superior parietal gyrus, angular gyrus, marginal gyrus, occipito-temporal gyrus, superior frontal gyrus, and occipital gyrus were also delineated (Fig. 6).

**Fig.6.**
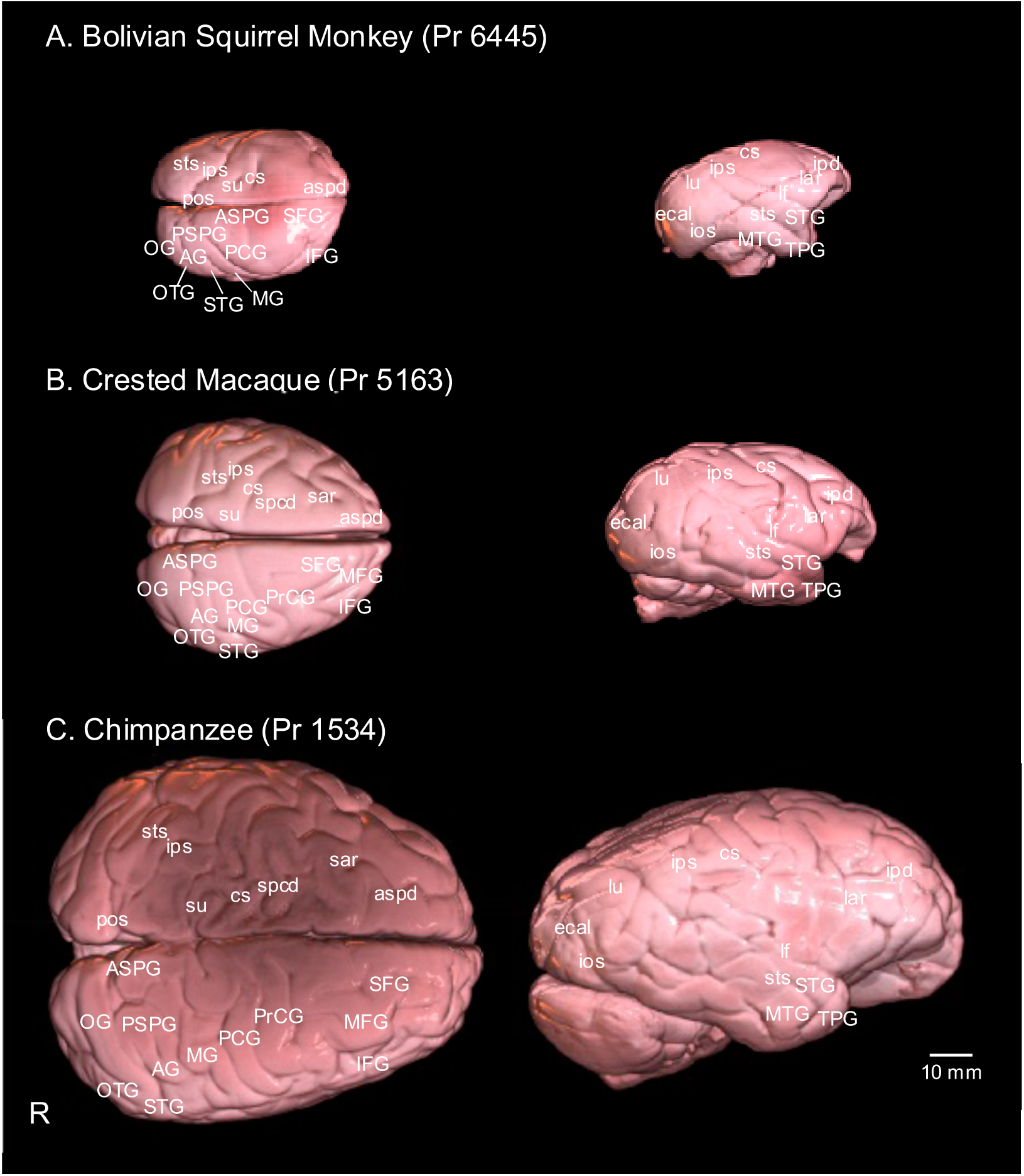
Demonstration of neuroanatomical details revealed in the brain surface. The image shows lateral (top) and dorsal (bottom) views of the brain surface from T2-weighted images in (A) Bolivian Squirrel Monkey (Pr 6445), (B) Crested Macaque (Pr 5163), and (C) Chimpanzee (Pr 1534). Neuroanatomical abbreviations in this MRI database are presented in accordance with the histological atlas of the rhesus monkey brain (Paxinos et al. 2000) and MRI and DTI atlas of the human brain (Mori et al., 2005; Oishi et al., 2010). Abbreviations of features in the brain surface: AG, angular gyrus; aspd, anterior supraprincipal dimple; ASPG, anterior superior parietal gyrus; CG, cingulate gyrus; cs, central sulcus; ecal, external calcarine sulcus; iar, inferior arcuate sulcus; IFG, inferior frontal gyrus; ios, inferior occipital sulcus; ipd, infraprincipal dimple; ips, intraparietal sulcus; lf, lateral fissure; lu, lunate sulcus; MFG, middle frontal gyrus; MG, marginal gyrus; MTG, middle temporal gyrus; OG, occipital gyrus; OTG, occipito-temporal gyrus; pos, parietooccipital sulcus; PrCG, precentral gyrus; PSPG, posterior superior parietal gyrus; sar, superior arcuate sulcus; SFG, superior frontal gyrus; spcd, superior precentral dimple; STG, superior temporal gyrus; sts, superior temporal sulcus; su, superior postcentral dimple; TPG, temporal polar gyrus.

T2-weighted contrast imaging enables detailed anatomical delineation in deep brain regions. For example, in chimpanzee (Pr1534), the T2-weighted image provided striking contrasts making it possible to delineate the accumbens, basal ganglia (putamen, external globus pallidus, internal globus pallidus), thalamus, mammillothalamic tract, periaqueductal gray, pulvinar, caudate nucleus, hippocampus, and superior colliculus from the mid-axial plane (Fig. 7). At the mid-coronal plane, the caudate nucleus, thalamic nucleus, and basal ganglia were identified (Fig. 8). Nearly adjacent to the mid-sagittal plane, 10 cerebellar lobules were distinctly delineated (Fig. 9).

**Fig.7.**
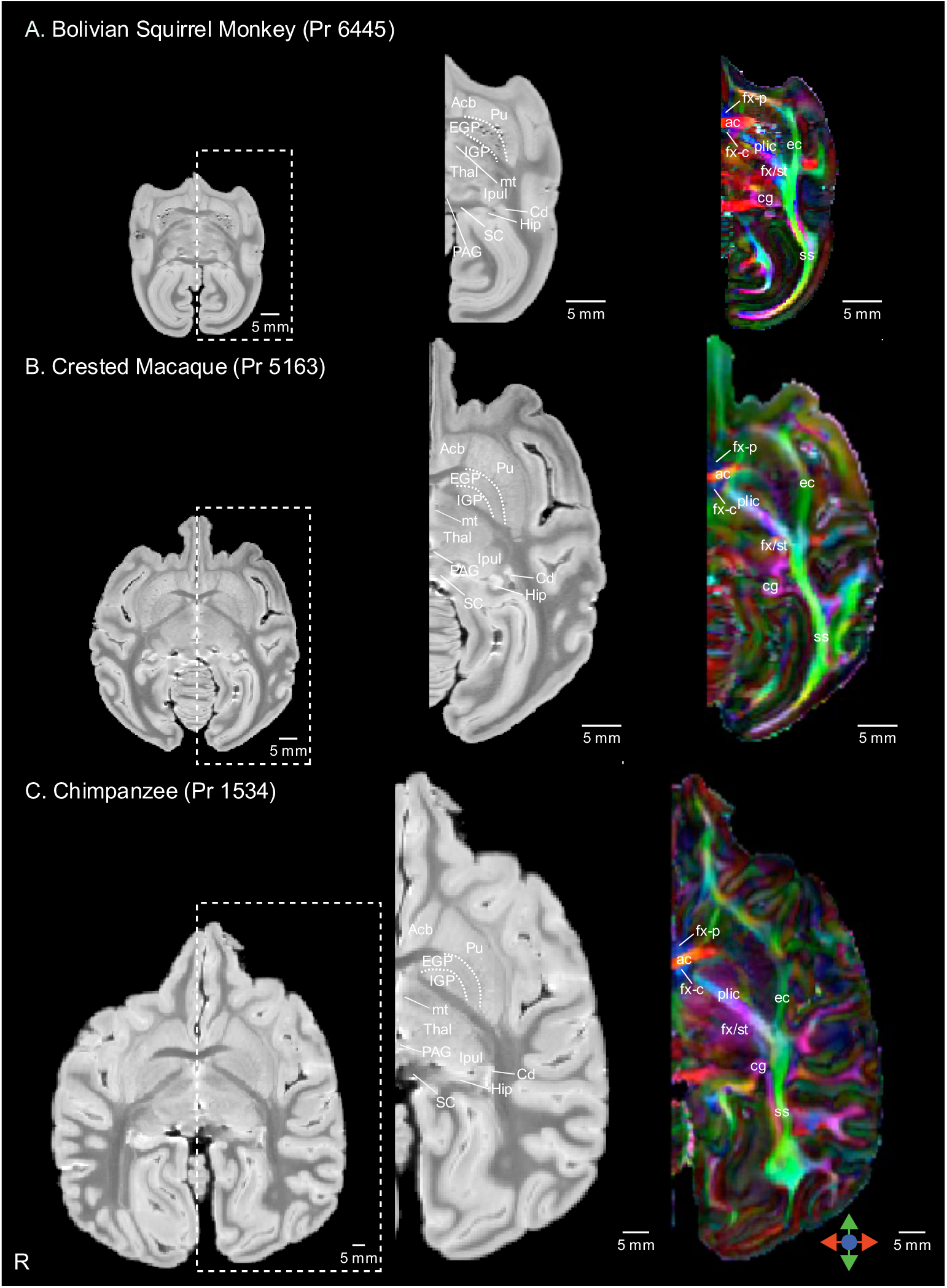
Demonstration of neuroanatomical details revealed with high-spatial-resolution T2-weighted image and DTI color map from mid-axial plane. The image shows T2-weighted images (middle column) and Color maps (right column) from the mid-axial plane in (A) Bolivian Squirrel Monkey (Pr 6445), (B) Crested Macaque (Pr 5163), and (C) Chimpanzee (Pr 1534). Red, green and blue indicate fiber orientations along medial–lateral, anterior–posterior, and superior–inferior axes, respectively. Neuroanatomical abbreviations in this MRI database are presented in accordance with the histological atlas of the rhesus monkey brain (Paxinos et al. 2000) and MRI and DTI atlas of the human brain (Mori et al., 2005; Oishi et al., 2010). Abbreviations in T2-weighted images: Acb, accumbens; Cd, caudate nucleus; EGP, external globus pallidus; Hip, hippocampus; IGP, internal globus pallidus; GN, geniculate nucleus; Ipul, inferior pulvinar; mlf, medial longitudinal fasciculus; PAG, periaqueductal gray; Pu, putamen; SC, superior colliculus; Thal, thalamus. Abbreviations in Color maps: ac, anterior commissure; cg, cingulum; ec, external capsule; fx, fornix; fx-c, column of the fornix; fx-p, precommissural part of the fornix; plic, posterior limb of the internal capsule; ss, sagittal stratum; st, stria terminalis.

**Fig.8.**
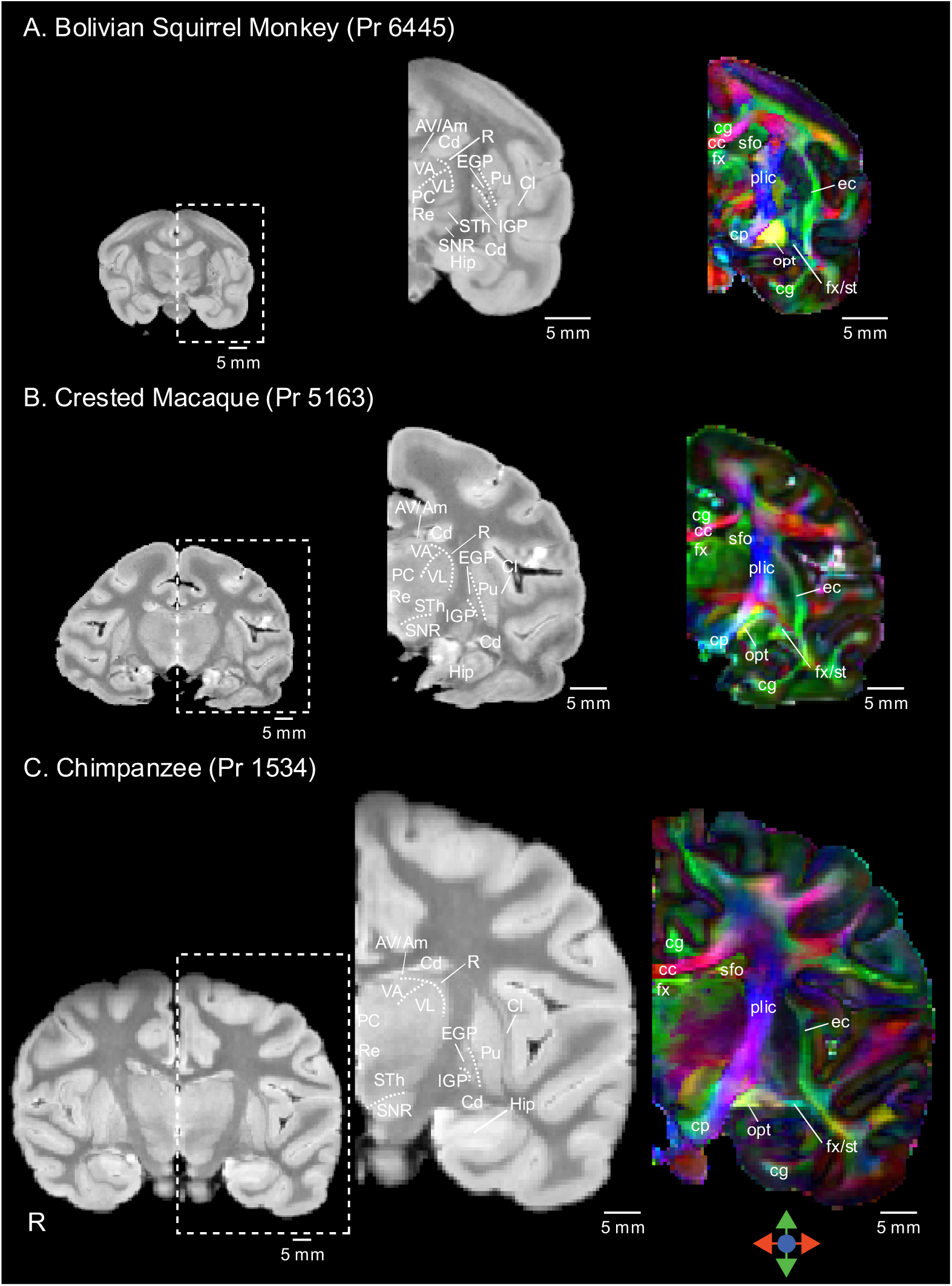
Demonstration of neuroanatomical details revealed with high-spatial-resolution T2-weighted image and DTI color map from mid-coronal plane. The image shows T2-weighted images (middle column) and Color maps (right column) from the mid-coronal plane in (A) Bolivian Squirrel Monkey (Pr 6445), (B) Crested Macaque (Pr 5163), and (C) Chimpanzee (Pr 1534). Red, green and blue indicate fiber orientations along medial–lateral, anterior–posterior, and superior–inferior axes, respectively. Neuroanatomical abbreviations in this MRI database are presented in accordance with the histological atlas of the rhesus monkey brain (Paxinos et al. 2000) and MRI and DTI atlas of the human brain (Mori et al., 2005; Oishi et al., 2010). Abbreviations in T2-weighted images: Am, anteromedial thalamic nucleus; AV, anteroventral thalamic nucleus; Cd, caudate nucleus; Cl, claustrum; EGP, external globus pallidus; GP, globus pallidus; Hip, hippocampus; IGP, internal globus pallidus; PC, paracentralis thalamic nucleus; Pu, putamen; R, reticularis thalamic nucleus; Re, reuniens thalamic nucleus; SNR, substantia nigra; STh, subthalamic nucleus; Thal, thalamus; VA, ventralis thalamic anterior nucleus; VL, ventralis thalamic lateralis nucleus. Abbreviations in Color maps: cc, corpus callosum; cg, cingulate; cp, cerebral peduncle; ec, external capsule; fx, fornix; opt, optic tract; plic, posterior limb of the internal capsule; sfo, superior fronto-occipital fasciculus; st, stria terminalis.

**Fig.9.**
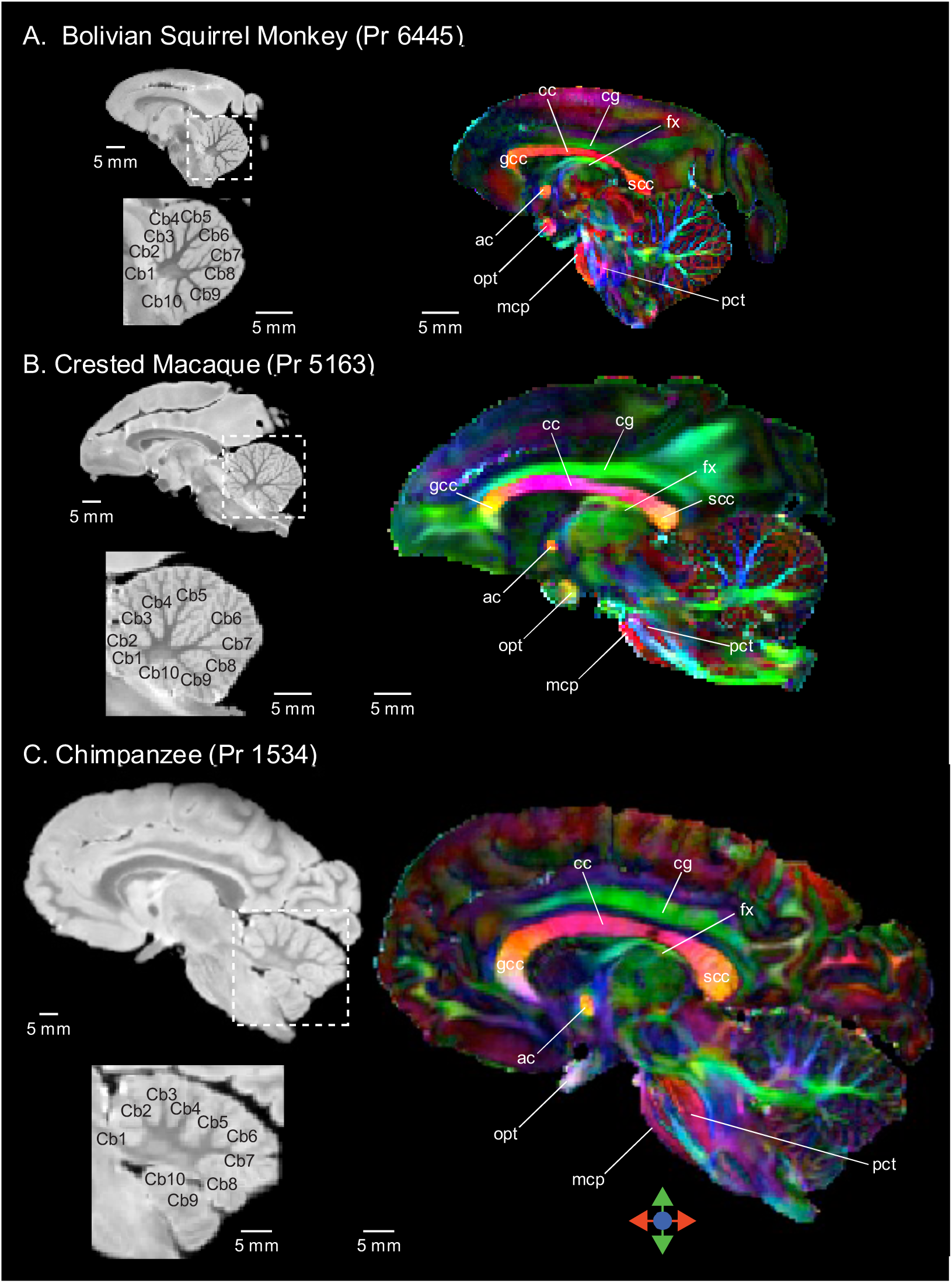
Demonstration of neuroanatomical details revealed with high-spatial-resolution T2-weighted image and DTI color map from almost mid-sagittal plane. The image shows T2-weighted images (middle column) and Color maps (right column) from almost mid-sagittal plane in (A) Bolivian Squirrel Monkey (Pr 6445), (B) Crested Macaque (Pr 5163), and (C) Chimpanzee (Pr 1534). Red, green and blue indicate fiber orientations along medial–lateral, anterior–posterior, and superior–inferior axes, respectively. Neuroanatomical abbreviations in this MRI database are presented in accordance with the histological atlas of the rhesus monkey brain (Paxinos et al. 2000) and MRI and DTI atlas of the human brain (Mori et al., 2005; Oishi et al., 2010). Abbreviations in T2-weighted images: Cb1, cerebellar lobule 1; Cb2, cerebellar lobule 2; Cb3, cerebellar lobule 3; Cb4, cerebellar lobule 4; Cb5, cerebellar lobule 5; Cb6, cerebellar lobule 6; Cb7, cerebellar lobule 7; Cb8, cerebellar lobule 8; Cb9, cerebellar lobule 9; Cb10, cerebellar lobule 10. Abbreviations in Color maps: ac, anterior commissure; cc, corpus callosum; cg, cingulum; dscp, decussation of the superior cerebellar peduncles; fx, fornix; gcc, genu of the corpus callosum; mcp, middle cerebellar peduncle; opt, optical tract; pct, pontocerebellar tract; scc, splenium of the corpus callosum.

### 3.2. Anatomical features and white matter bundles from diffusion tensor images

Representative scalar maps derived from DTIs are shown in Figs. 7, 8, and 9. Structural orientation-based contrast in the color maps enabled us to delineate well-aligned structures, such as white matter bundles and columnar structures within the cortex. For example, in Bolivian squirrel monkey (Pr1534), crested macaque (Pr5163), and chimpanzees (Pr1534), the respective color maps provided striking contrasts allowing us to delineate the anterior commissure, external capsule, posterior limb of the internal capsule, sagittal stratum, column of the fornix, precommissural part of the fornix, fornix, stria terminalis, and cingulum at the mid-axial plane (Fig. 6). At the mid-coronal plane, the posterior limb of the internal capsule, external capsule, cerebral peduncle, optic tract, corpus callosum, cingulate, fornix, superior fronto-occipital fasciculus, and stria terminalis were identified in the three species (Fig. 7). Close to the mid-sagittal plane, the anterior commissure, optical tract, pontocerebellar tract, corpus callosum, genu of the corpus callosum, splenium of the corpus callosum, cingulum, fornix, and middle cerebellar peduncle were distinctly delineated among the three species (Fig. 8). Using the DTIs, we also succeeded in constructing white matter fiber bundles three-dimensionally by tractography (Fig. 10).

**Fig.10.**
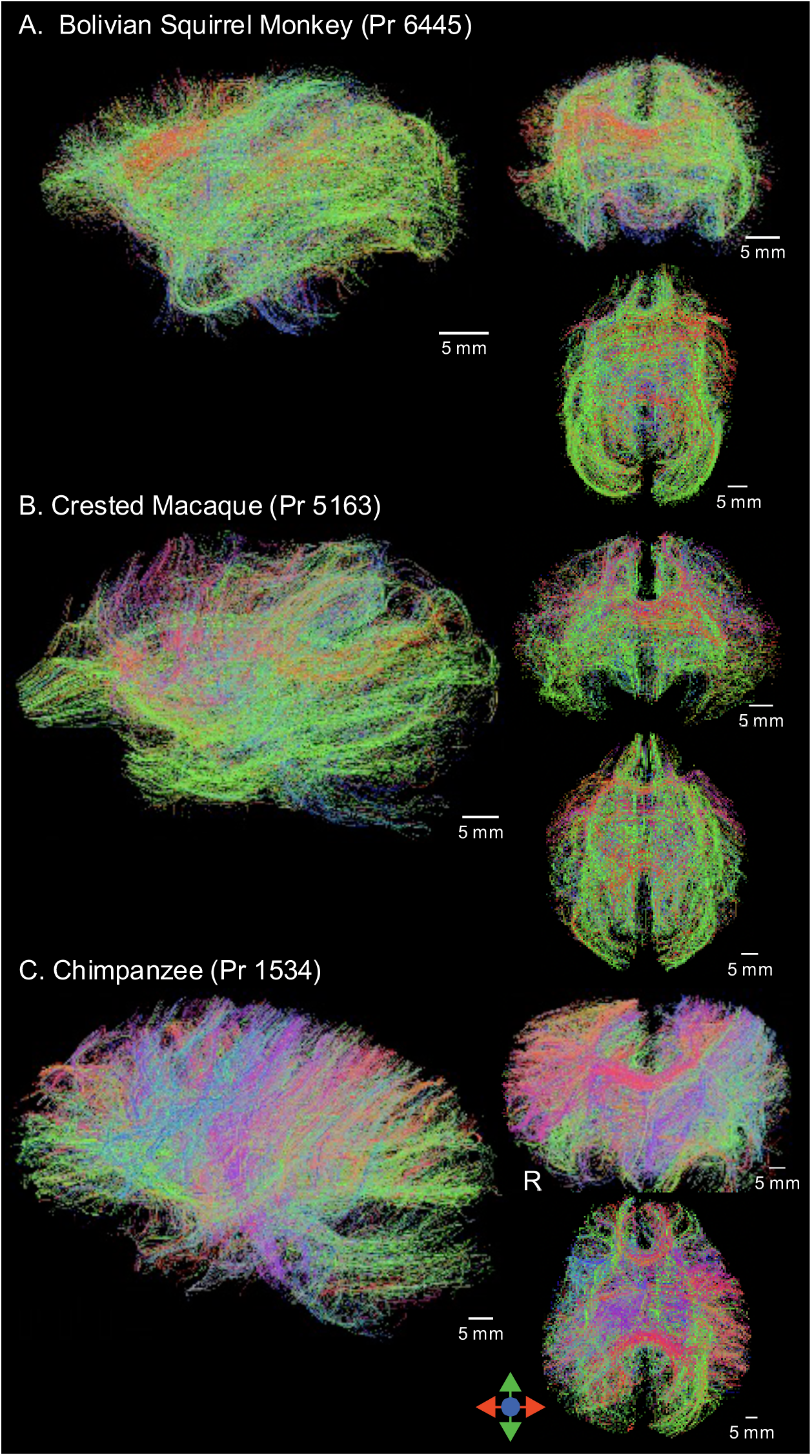
Demonstration of whole-brain tractography from DTIs. The image shows whole-brain tractography from DWIs in (A) Bolivian Squirrel Monkey (Pr 6445), (B) Crested Macaque (Pr 5163), and (C) Chimpanzee (Pr 1534). Red, green and blue indicate fiber orientations along medial–lateral, anterior–posterior, and superior–inferior axes, respectively.

### 3.3. Incidental findings

The brain sample of the pig-tailed macaque (Pr3010) included a space-occupying lesion that was observed around the left temporal area in the T2-weighted images (Fig.11). A low signal was observed in the same area in the color map (Fig.11). At the macroscopic anatomical level, this lesion was not visible from the brain surface and therefore was not recognized until the brain MRI scan was performed. Although the reported gross anatomical observations of the monkeys included swelling of the hilum of the lungs, lymph nodes, and spleen, this brain lesion was not visible from the brain surface, and therefore was not recognized until the brain MRI scan was performed.

**Fig.11.**
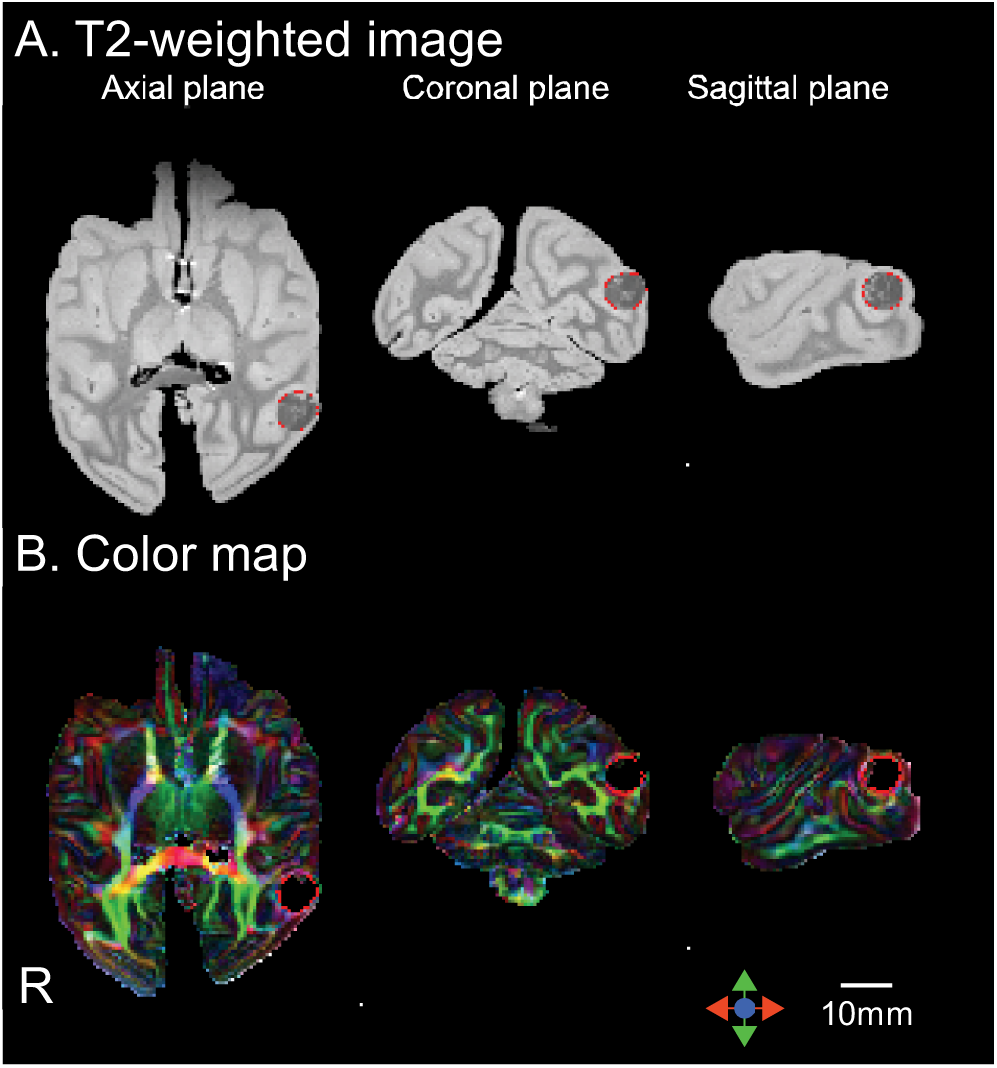
Images of brain with space-occupying lesion. The images show T2-weighted images, DWIs, and Color maps in a pig-tailed macaque (Pr3010). (A) T2-weighted images and (B) Color maps. Axial, coronal, and sagittal planes are indicated. Areas surrounded by red lines represent the space-occupying lesion. Red, green and blue indicate fiber orientations along medial–lateral, anterior– posterior, and superior–inferior axes, respectively.

## 4. Discussion

Toward our long-term goal of establishing the JMC Primates Brain Imaging Repository, which will include images from more than 100 nonhuman primate species for harmonious facilitation of the fields of comparative neuroscience, animal ethics, animal welfare and conservation, MRIs from 12 individuals belonging to nine nonhuman primate species were released in the second phase. Combining these data with those from the first phase, we obtained MRI data from 28 brain samples of 20 species in total. Thus, we developed the first nonhuman primate brain image repository from 20 different nonhuman primate species with a consistent scan protocol using 9.4-T high-resolution MRI. Our study successfully acquired both high-resolution structural MRIs and DTIs of nonhuman primate whole brains, and we will release these data to the research community in the near future.

### 4.1. Technical improvements from first phase in high-resolution postmortem MRI

In the present phase, we made the following three improvements over the imaging performed in the first phase (Sakai et al., 2018).

#### 4.1.1. Technical improvement of scanning of large postmortem brains

We succeeded in scanning whole large postmortem brains, such as brains of chimpanzees, with high-resolution. There was a brain size limitation in the first phase (Sakai et al., 2018), namely, that it was not possible to image brain samples with a maximum diameter of approximately 70 mm or larger due to the bore diameter limitation of the 9.4-T MRI equipment (approximately 200 mm diameter) (Sakai et al., 2018). In the present study, we were able to scan relatively large brain samples, such as chimpanzee brain samples, using a 9.4-T MRI machine with a wider bore (approximately 300 mm diameter).

#### 4.1.2. Technical improvement to reduce image artifacts and increase signal-noise ratio

In the first phase, deaeration before scanning was not performed, and some brain images had image artifacts or insufficient single-noise ratio (SNR) (Sakai et al., 2018). These problems were caused by air bubbles from brain samples and the container that severely decreased the quality of *ex vivo* MRI data, especially *ex vivo* DTIs. In the second phase, we succeeded in completely eliminating air bubbles from both the brain sample and the container by deaeration with a vacuum pump before scanning.

In the first phase, we also found that some brain images of large brain samples had concentric contrast inhomogeneity (Sakai et al., 2018), but it is difficult to explain the exact cause of this artifact because gadopentetate dimeglumine (GD) usually enhances image contrast. It is standard practice to immerse the fixed brain samples in GD-PBS for more than 1 week prior to *ex vivo* MRI scanning of mouse brain samples (Johnson et al., 2002). Although this preprocessing was also used for the *ex vivo* MRI scanning of marmoset brain samples (Hikishima et al., 2013), we found that it cannot be used for scanning of the samples of brains larger than the marmoset brain.

In the present study, we were able to avoid problems with brain image contrast by scanning the nonhuman primate brain specimens without GD-PBS. The concentric contrast inhomogeneity appears to be due to nonuniform penetration of GD-PBS into the brain samples, but providing the exact cause of this artifact is difficult because GD usually enhances image artifacts. Further investigation will be needed to determine the mechanism that generates this imaging artifact, but nonhuman primate brain samples larger than marmoset brain specimens may need to be scanned without GD-PBS.

#### 4.1.3. Points to consider for cross-species comparison study using this image repository

We made some technical improvements in MRI scanning, as described above, in the second phase. Users of this image repository need to pay attention to the differences in scanning methods when performing cross-species comparison analysis by combining the nonhuman primate brain MRIs from the first and the second phases. In the future, we plan to re-scan the samples analyzed in the first phase with the improved scanning methods.

### 4.2. Contributions from Japan Monkey Centre Primates Brain Imaging Repository

We expect four significant contributions from our repository, namely, contributions to comparative neuroscience, animal ethics, animal welfare, and conservation.

#### 4.2.1. Contribution to comparative neuroscience

This nonhuman primate imaging database provides a resource for comparative neuroscience to investigate the brain morphology and anatomical connectivity of each species. We anticipate that this imaging repository of T2 images and DTIs will facilitate discoveries not only in the field of brain evolution but also in the field of connectomics, a technique that builds and studies comprehensive maps of the complex network of connections within the brain.

Furthermore, a variety of information about the species whose brains we scanned, such as gestational period, lifespan, allomothering and social structure, as well as individual data such as body weight and brain weight, are provided. For example, users of this database can investigate the relationship between the encephalization quotient (EQ), EQ = Ea/Ee, which indicates the extent to which the brain size of a given species, Ea, deviates from the expected brain size, Ee (Jerison, 1973), and the absolute or relative size of the cerebral cortex to understand the remarkable encephalization during human evolution.

The users also can analyze the relationship between social scores and the absolute or relative size of the brain regions or connection strengths of the white matter bundles to investigate the social abilities among primates.

#### 4.2.2. Contribution to animal ethics and animal welfare

The repository provides opportunities to optimize protocols for postmortem brain scans. We applied state-of-the-art scanning protocols to acquire the *ex vivo* T2 images and DTIs of nonhuman primate brains using high-magnetic-field MRI equipment. Information concerning the scanners, scan preparation, and scan protocols is fully open. This provides opportunities for MR physicists to develop novel MRI sequences optimized for primate brain samples of various sizes, or for researchers who are interested in image post-processing to develop novel image analysis methods. Our imaging database will serve as a resource for such new scan protocols or analysis methods in nonhuman primate imaging studies.

Sharing of such high-resolution brain images and scan information is very important from the perspective of animal ethics, as it enables the development of appropriate scan protocols for *in vivo* and *ex vivo* imaging and a reduction of the number of individuals needed for statistical comparisons in nonhuman primate brain imaging studies at different research facilities around the world. The nonhuman brain images of our repository can be analyzed in various ways without limitations. In addition, these images will allow researchers who are not affiliated with laboratories possessing nonhuman primate models to participate in nonhuman primate brain imaging.

This repository can potentially serve as a disease reference for nonhuman primates. In the second data series, a space-occupying lesion was found incidentally in the left temporal area of a pig-tailed macaque (Pr3010), and a histopathological examination will be performed to identify the etiology of this lesion. Once a diagnosis is established, the image can be used as a reference file for veterinary education and practice.

There are many neuroradiology reference files or teaching files, both in printed and electronic forms, for human brain diseases (Poldrack and Gorgolewski, 2014). However, only a few veterinary neuroradiology reference files for nonhuman primates are currently available. Some examples are case studies of a lateral ventriculomegaly and a mild hippocampal atrophy in a young marmoset (Sadoun et al., 2015), a subacute necrotizing encephalopathy in a pig-tailed macaque (Bielefeldt-Ohmann et al., 2004), an ependymal cyst in a cynomolgus macaque (*Macaca fascicularis*) (Bergin et al., 2008), a neurocysticercosis in a rhesus macaque (*Macaca mulatta*) (Johnston et al., 2016), and intracranial arachnoid cysts in a chimpanzee (Kaneko et al., 2013; Miyabe-Nishiwaki et al., 2014).

In order to develop neuroradiology reference files or educational files for nonhuman primate brain diseases, it will be necessary to accumulate more electronic images from such cases. We expect that our repository will contribute to accelerating the development of veterinary neuroradiology for nonhuman primates.

#### 4.2.3. Contribution to conservation

The JMC Primates Brain Imaging Repository will be a permanent repository of primate brains, since the images are stored as digital data. Unlike fixed brains, digital data are not subject to deterioration, and they can be distributed world-wide. This repository is especially important for preserving information regarding endangered species, since this precious information would otherwise be lost. We will continue to acquire additional whole brain MRIs from various endangered nonhuman primates such as barbary macaques, orangutans and gorillas in the near future.

### 4.3. Limitations

Although significant technical improvements have been made since the first phase, the second phase dataset still has the following limitations.

#### 4.3.1 Postmortem brain samples

The accurate times of death of the JMC’s brain samples are not necessarily clear, as the primates often died spontaneously. Small artifactual incisions were at times made during brain extraction from the cranium despite the best efforts of skilled veterinarians to avoid such artifacts.

#### 4.3.2 Brain sample shrinkage after formalin fixation

Users of this database should take into account shrinkage of the brain and changes of the brain morphology after formalin fixation. Average shrinkage of formalin-fixed brain is approximately 10–20% (de Guzman et al., 2016; Kinoshita et al., 2001; Quester and Schröder, 1997). The degree of shrinkage also depends on the water content of brain tissue. Especially, the degree of shrinkage of non-myelinated immature brain samples, which contain water-rich tissue, would be greater than that of mature brain samples (Kinoshita et al., 2001). Our database includes immature brain samples, such as from a chimpanzee (Pr1534, young). The shrinkage rates should therefore be considered when analyzing the size, shape, and connection strengths of white matter bundles of the brain in formalin-fixed T2 images and DTIs of brains, and especially in non-myelinated brains.

#### 4.3.3 *Ex vivo* diffusion tensor images of nonhuman primate brains as experimental data

Similarly to previous *ex vivo* DTIs (details in (Oishi et al., 2020)), there are several issues related to *ex vivo* DTIs in the second phase as well. Low temperature and tissue fixation are known to reduce the diffusion coefficient (Schmierer et al., 2008), mandating the use of a high b value to sensitively detect diffusion, although increasing b values and resolutions are concomitant with a decrease in SNR. A high magnetic field was required to obtain DTI with an SNR high enough to delineate detailed anatomical structures on a micro and macro scale. DTI-derived scalar values are affected by scan parameters and magnetic fields (Chung et al., 2013), as well as by tissue deformation occurring during excision and fixation (Guilfoyle et al., 2003; Sun et al., 2005). Therefore, users should take care to compare the FA and ADC values in this repository to those obtained from other *ex vivo* or *in vivo* DTI studies.

## 5. Conclusion

The JMC Primates Brain Imaging Repository is an *ex vivo* MRI database of nonhuman primate brains that can facilitate scientific discoveries in the field of comparative neuroscience. Our repository can also promote animal ethics and welfare in experiments with nonhuman primate models by optimizing methods for *in vivo* and *ex vivo* MRI scanning of brains and supporting veterinary neuroradiological education. The repository is important for conservation of primate brain information, preserving data of the brains of various primates, including endangered species, in a permanent digital form.

As the second phase, we are opening a digital collection of structural MRIs and DTIs obtained from 9 primate species to the research community. Its use by scientists belonging to research fields outside traditional neuroscience and primatology, such as computer science, mathematical modeling, and medicine is strongly encouraged. It may be a long road requiring considerable time and effort, but we will continue to work together to develop this repository in order to achieve the goal of collecting and publishing brain MRIs of more than 100 primate species.

## 6. Access to the Japan Monkey Centre Primates Brain Imaging Repository

This publication coincides with the public release of the JMC Primates Brain Imaging Repository. The images can be browsed online from the following JMC website (http://www.j-monkey.jp/BIR/index_e.html). Users can access the images and non-image information such as cause of death and autopsy findings that were released in the first and second phase of this repository after completion of a collaborative research agreement with representative authors of this article.

## Acknowledgments

We thank N. Kimura for advice on primate brain samples, and the JMC for permitting us to use the JMC primate brain sample collection. We also thank Drs. E. Nakajima and A. Gerz for help with manuscript editing.

## Funding

This work was financially supported by JSPS KAKENHI Grant for Young Scientists (B) (#26870827 and #17K18097 to T.S.), a JSPS Postdoctoral Fellowship for Research Abroad (#490 to T.S.), Brain/MINDS (JP17dm0207002 to T.S., JP20dm0207001 to H.O./J.H.) and Brain/MINDS Beyond (JP20dm0307007h0002 to T.H./T.M and JP20bm0307005 to N.S.) by AMED, the Fakhri Rad Brite Star award from the Department of Radiology, Johns Hopkins University School of Medicine (to K.O.), and JSPS KAKENHI Grant for Scientific Research (JP20H03630 to J.H.). This work was also supported by the JMC collaborative research program (#2014013, #2015019, and #2016017 to T.S.) and the Cooperative Research Program of the Primate Research Institute, Kyoto University (#H25-E31 and# H26-C7 to H.O, and #H27-D23 to T.S.).

## Notes

### Competing Interest Statement

The authors have declared no competing interest.

